# Learning sculpts orthogonal task manifolds for continual skill learning in recurrent networks

**DOI:** 10.64898/2026.02.15.705283

**Authors:** Zihan Liu, Anno Kurth, Yuma Osako, Toshitake Asabuki

## Abstract

Humans and animals can learn and seamlessly perform a vast repertoire of behaviors. However, how neural populations incorporate new skills without disrupting previously learned ones remains poorly understood. This challenge is known as catastrophic forgetting in artificial neural networks and is especially severe in recurrent networks, where computation relies on stable internal dynamics. While current machine learning approaches often rely on explicit weight-protection strategies, such approaches do not directly address this challenge. Here we show that orthogonal task manifolds can emerge in recurrent neural networks from a local predictive, error-driven learning rule. Our model preserves task-specific latent dynamics that remain resilient to interference from subsequent learning. Using low-rank connectivity ablations, we causally isolate these latent dynamics, selectively impairing individual tasks while leaving others intact. We further show that the proposed principle generalizes beyond low-dimensional to high-dimensional naturalistic movie replay, suggesting a scalable mechanism for continual learning. Our results identify a solution to catastrophic forgetting by preserving previously learned dynamics in recurrent connectivity, thereby providing a mechanistic bridge between artificial recurrent networks and biological neural circuits.

## Introduction

The ability to continuously acquire new skills without overwriting previously learned ones is a hallmark of biological intelligence. In contrast, when artificial neural networks are trained sequentially on multiple tasks (continual learning), they often suffer from catastrophic forgetting, whereby performance on earlier tasks collapses as new information is incorporated (McCloskey & Cohen, 1989; Ratcliff, 1990; French, 1999). This limitation poses a major obstacle to continual learning in machines and raises a fundamental question: how do biological circuits preserve previously acquired computations while remaining plastic enough to learn new ones without catastrophic interference?

A variety of strategies have been developed to mitigate forgetting in feedforward networks. Weight-consolidation methods estimate parameter importance and penalize deviations during subsequent learning (Kirkpatrick et al., 2017; Zenke et al., 2017; Aljundi et al., 2018), while replay-based strategies interleave past experience during new learning to stabilize performance (Robins, 1995; Shin et al., 2017). Other approaches reduce interference by allocating task-specific resources, including modular architectures and progressive expansion (Rusu et al., 2016), or by explicitly constraining weight updates, for example using gradient projection or orthogonalization (Lopez-Paz & Ranzato, 2017; Zeng et al., 2019; Farajtabar et al., 2020). Despite their practical success, these methods primarily target static input-output mappings and do not directly address continual learning in recurrent neural networks (RNNs), where computation is inextricably tied to self-generated internal dynamics (Sussillo & Abbott, 2009; Sussillo & Barak, 2013).

Continual learning in RNNs is uniquely challenging because task representation is embedded not only in synaptic weights but also in the geometry of population trajectories produced by recurrent interactions (Sompolinsky et al., 1988; Rabinovich et al., 2006; Shenoy et al., 2013). Interference can therefore occur through two coupled modes. First, synaptic updates can overwrite previously learned recurrent dynamics (Fu et al., 2012; Nishiyama & Yasuda, 2015; Wright et al., 2025). Second, even modest perturbations of recurrent connectivity can reshape latent trajectories (Finkelstein et al., 2026), disrupting temporal computations that rely on persistent internal dynamics. Understanding how recurrent circuits avoid such interference is particularly relevant to neuroscience, because recurrent dynamics are thought to underlie working memory, decision-making, and motor control (Mante et al., 2013; Sussillo et al., 2015).

A growing body of experimental and modeling work suggests that recurrent computations are organized within low-dimensional manifolds of population activity (Cunningham & Yu, 2014; Gallego et al., 2017; Saxena & Cunningham, 2019). Such manifolds have been identified across diverse brain regions and behaviors, including motor cortex during movement generation (Churchland et al., 2012; Gallego et al., 2018), prefrontal cortex during context-dependent decision making (Mante et al., 2013), and navigation-related circuits (Harvey et al., 2012; Gardner et al., 2022). Manifold geometry is increasingly linked to multi-task computation, where context can be implemented through rotations or gating of neural subspaces rather than wholesale circuit rewiring (Mante et al., 2013; Yang et al., 2019; Panichello & Buschman, 2021; Naumann et al., 2022; Kim et al., 2025; Osako et al., 2025). Theoretical studies further suggest that allocating tasks to orthogonal or minimally overlapping dynamical subspaces can, in principle, eliminate catastrophic forgetting in recurrent systems (Duncker et al., 2020). However, existing implementations often rely on explicit architectural constraints (Rusu et al., 2016), global optimization objectives (Duncker et al., 2020), or offline procedures that require simultaneous access to information from multiple tasks (Kirkpatrick et al., 2017; Zenke et al., 2017; Aljundi et al., 2018), making them incompatible with continual learning through local synaptic updates. How such task-specific manifold organization can emerge incrementally through local synaptic plasticity, without access to past task information, remains an open question.

Here, we show that continual learning in RNNs can emerge from modulating the feedback signals that drive synaptic plasticity. Using a local predictive plasticity rule (Asabuki and Clopath, 2025), we train networks on sequential stimulus-response mappings while switching only the feedback pathways that govern recurrent weight updates, without providing task-identity input to recurrent units. We find that distinct feedback vectors are sufficient to guide population trajectories and synaptic updates into task-specific subspaces. This separation of task manifolds stabilizes task-specific latent dynamics against subsequent interference, enabling both long-term retention and accelerated re-learning through feedback-specific reactivation. We causally isolate these task-specific modes using targeted low-rank connectivity ablations, and we demonstrate the generality of the principle by extending it to high-dimensional naturalistic movie replay. Together, our results identify feedback-driven manifold separation as a mechanistic and scalable route to continual learning in recurrent circuits.

## Results

### Feedback-driven predictive plasticity organizes task-specific manifolds

To understand how recurrent neural networks (RNNs) can learn multiple tasks without catastrophic forgetting, we first considered the geometric requirements for stable continual learning. In RNNs, task-specific computations are often embedded in population dynamics confined to effectively low-dimensional manifolds. We reasoned that to avoid interference, the network might separate task-relevant neural trajectories into distinct, minimally overlapping subspaces (Fig. 1A). In the following, we refer to the use of these subspaces as “manifold switching.”

**Figure 1.**
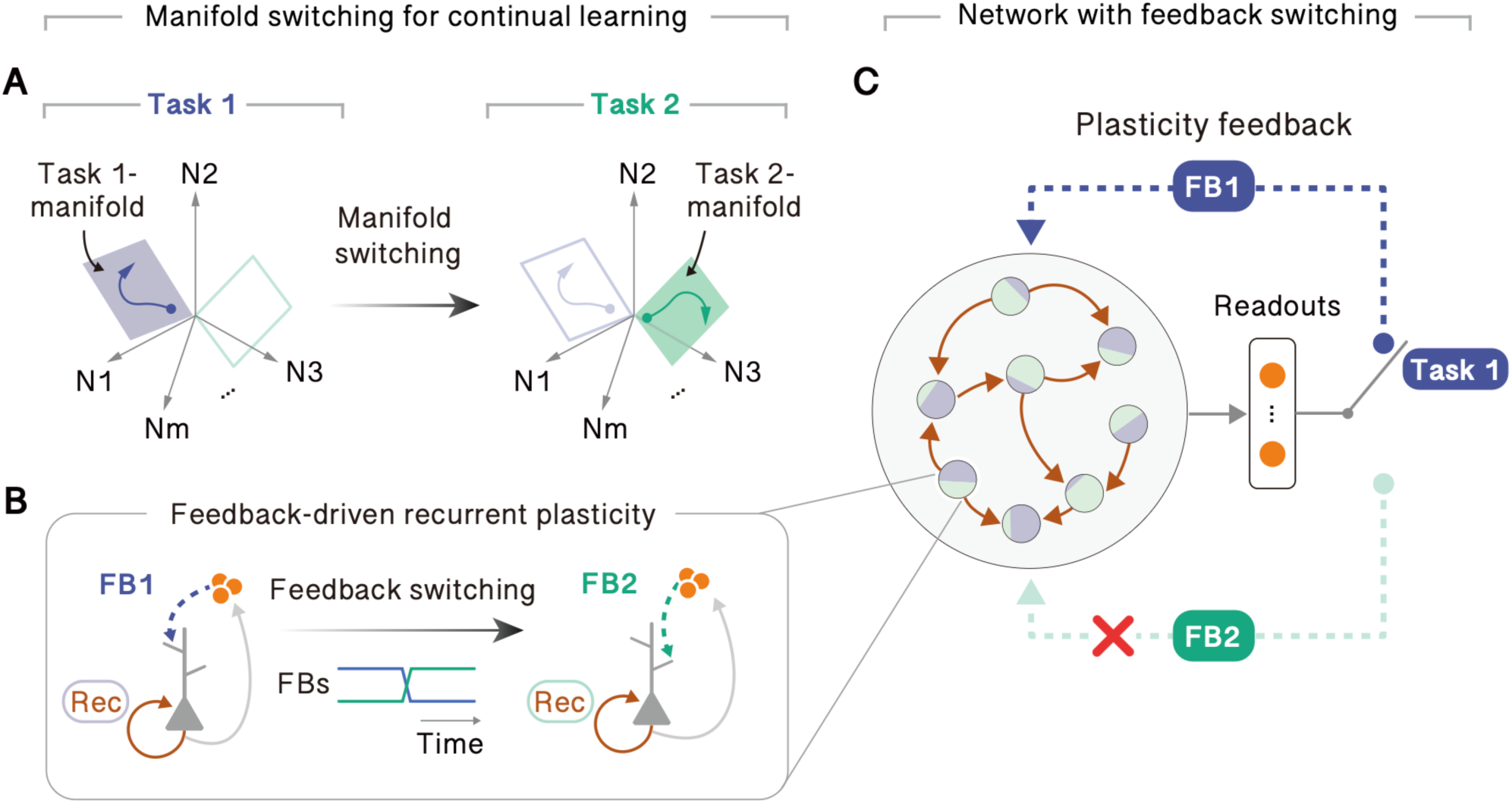
Feedback-driven switching of task manifolds in recurrent neural networks. **A** Task-specific population trajectories are confined to distinct, minimally overlapping manifolds, enabling switching between tasks while preserving previously learned dynamics. **B** Switching the active feedback pathway (from FB1 to FB2) provides a new target direction for synaptic plasticity, directing recurrent updates to embed task-specific computations into separate subspaces. **C** The network consists of a recurrently connected population where plasticity is governed by a local feedback-predictive rule. Crucially, the recurrent units receive no explicit task-identity inputs. Instead, task context is provided solely by changing the plasticity feedback pathways (FB1 and FB2) that drive recurrent weight updates.

We hypothesized that such geometric organization can be achieved through a feedback-driven predictive learning principle (Asabuki and Clopath, 2025). In this framework, task-specific feedback signals align population dynamics along the corresponding feedback vector via recurrent synaptic plasticity (Fig. 1B). Switching the task-specific feedback pathway therefore presents a new target direction, driving the plasticity rule to embed the new task’s dynamics in a distinct subspace.

To test this idea, we implemented this learning rule in an RNN model (Fig. 1C). Crucially, task context was manipulated solely by switching between randomly generated, high-dimensional feedback vectors for recurrent plasticity (FB1 and FB2), which were nearly orthogonal, while recurrent units received no explicit task-identity input. We show that feedback switching is sufficient to segregate task trajectories into minimally overlapping subspaces, preserving task-specific latent dynamics across task switches. For clarity, in most simulations, we externally switch the feedback signals across tasks to isolate the effect of feedback-specific plasticity. In principle, however, the task-specific feedback pathway can be generated autonomously by the network, and we later demonstrate one such mechanism.

### Task-specific feedback enables rapid memory reactivation without interference

To test whether feedback-driven plasticity protects previously learned dynamics from interference, we trained networks on a context-dependent binary choice task (Kim et al., 2025) in which the stimulus–response contingency reversed across two tasks (Fig. 2A). In Task 1, stimulus 1 cued a right response while stimulus 2 cued left; in Task 2, this contingency was reversed. Networks received one of two constant hold inputs during a delay period, followed by removal of that input to generate a binary motor output (Fig. 2B). We employed a three-phase training protocol: initial learning of Task 1 with feedback FB1 followed by learning of Task 2 with feedback FB2 and finally re-learning of Task 1 using either FB1 or FB2 (Fig. 2C). Network activity was initialized at the beginning of each trial to a predefined state (Methods). During the initial learning phases, error decreased monotonically within each task but increased abruptly at task transitions.

**Figure 2.**
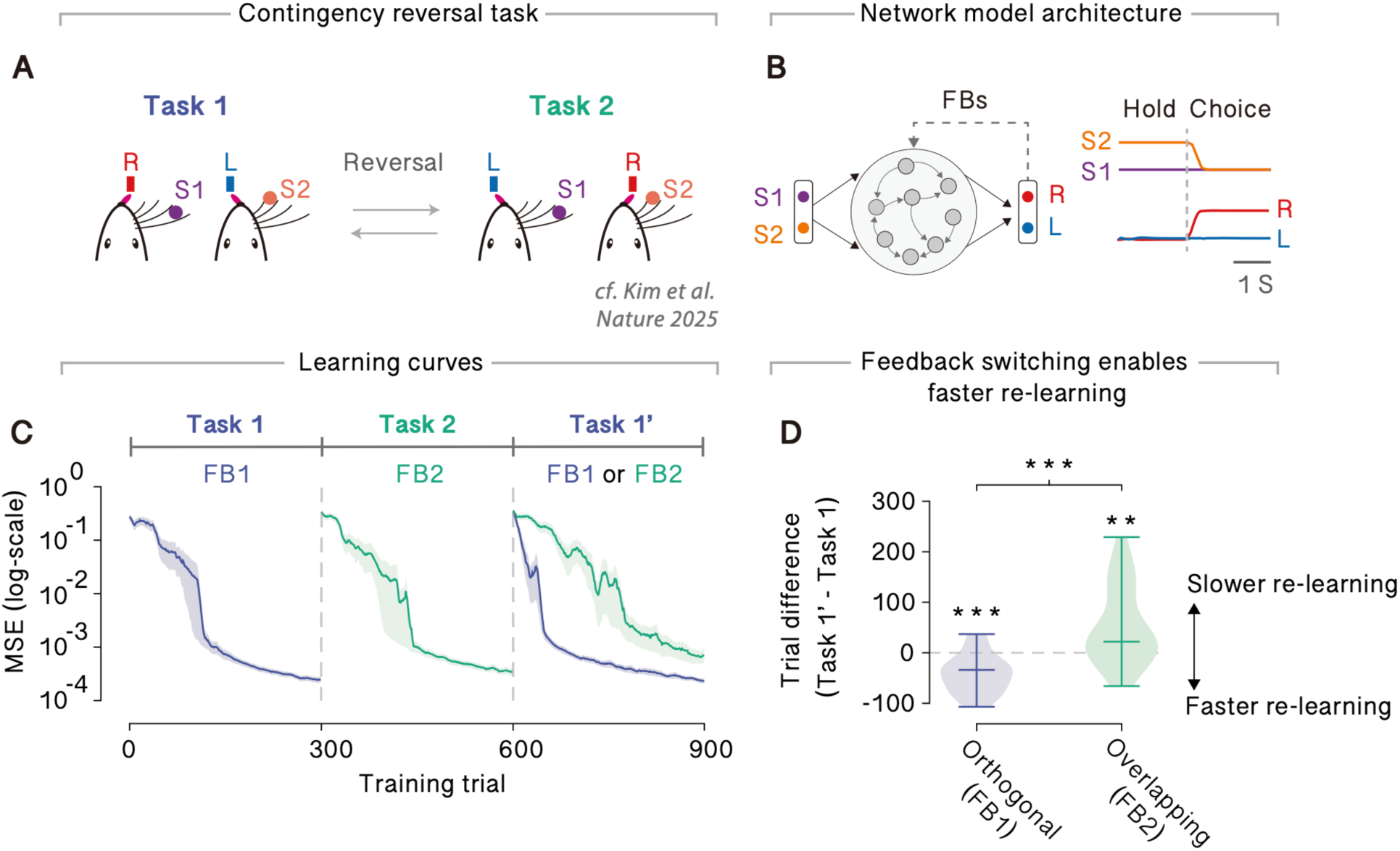
Switching of feedback signals reduces task interference. **A** Context-dependent binary choice task. In Task 1, stimulus 1 cues a right choice and stimulus 2 cues a left choice; this stimulus-choice contingency is reversed in Task 2 (Kim et al., 2025). **B** Schematic of the recurrent network model. The network receives one of two stimuli during the hold period and generates a binary choice output during the subsequent choice period. **C** Learning curves across three training phases: Task 1 learning with FB1, Task 2 learning with FB2, and Task 1 re-learning under different feedback conditions. Shaded areas represent s.d. over 20 trials. **D** Difference in the number of trials required to reach a mean squared error (MSE) threshold (MSE < 10⁻^3^) during Task 1 re-learning, relative to a novel-feedback baseline. In D, P values were obtained from two-sided Welch’s t-tests across 20 independent simulations (**p < 0.01; ***p < 0.001).

The critical test occurred during the re-learning phase of Task 1 (Fig. 2C). We compared re-learning under the original feedback FB1 (orthogonal) to re-learning under the misaligned feedback FB2 (overlapping). Re-learning was substantially faster with FB1 than with FB2, indicating selective reactivation of task-specific dynamics by aligned feedback. Consistent with this interpretation, the same dependence on feedback signal was observed in errors of the recurrent plasticity, accompanied by distinct reorganization of recurrent connectivity (Supplementary Fig. 1).

To quantify whether this effect reflected facilitation or interference relative to baseline learning, we compared re-learning speeds to a novel-feedback condition. Re-learning with FB1 was significantly faster than with novel feedback, indicating reuse of task-specific dynamics previously acquired during the initial training, whereas re-learning with FB2 was slower than novel feedback, indicating interference (Fig. 2D). Consistent with this result, preserving Task 1 dynamics also depended on the feedback used during Task 2 learning: using task-aligned feedback during Task 2 reduced interference and facilitated subsequent Task 1 re-learning (Supplementary Fig. 2).

Altogether, these results demonstrate that re-learning speed depends critically on feedback direction: task-aligned feedback enables rapid memory reactivation, whereas misaligned feedback fails to elicit such reactivation.

### Orthogonal feedback guides the segregation of task-specific neural manifolds

We then asked whether switching feedback pathways indeed induces switching of task manifolds. To this end, we examined the alignment between feedback signals and low-dimensional population activity during learning over the three task training phases shown above (Methods). Population activity was projected onto the first two principal components (PCs), which defined the dominant low-dimensional manifold for each phase. During Task 1 learning and re-learning, FB1 was strongly aligned with the dominant PCs, whereas FB2 showed little alignment (Fig. 3A). In contrast, during Task 2 learning, alignment shifted such that FB2 aligned with the dominant PCs while FB1 did not. Notably, this separation was not accompanied by changes in manifold dimensionality, which remained comparable across the two conditions (Supplementary Fig. 3). Indeed, during re-learning, population activity selectively reactivated the task manifold associated with the currently active feedback signal, remaining orthogonal to the other task manifold (Supplementary Fig. 4).

**Figure 3.**
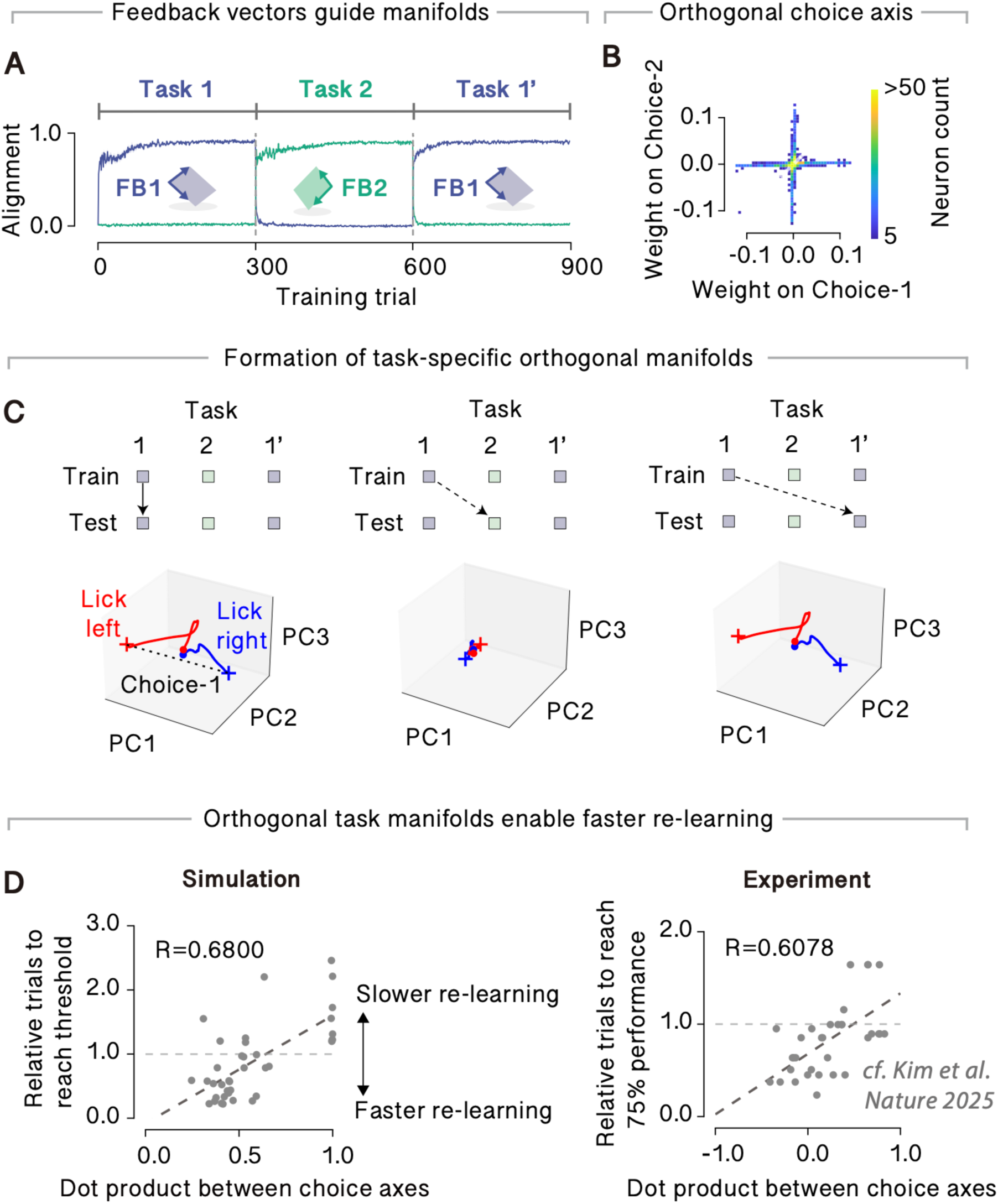
Task memories are encoded in separable neural manifolds. **A** Alignment between feedback signals and low-dimensional population activity across learning phases. During Task 1 learning and re-learning, FB1 aligned with the dominant population components, whereas FB2 showed little alignment. During Task 2 learning, alignment shifted such that FB2 aligned with the dominant components. Shaded areas represent s.d. over 20 trials. **B** Orthogonalization of task-relevant choice representations. Choice axes (e.g., Choice-1) separating left and right responses were approximately orthogonal across the two tasks. **C** Context-dependent engagement of task-specific manifolds. When population activity was projected onto the Task 1 subspace, left and right trajectories were clearly separated during Task 1 learning and re-learning, whereas trajectories from Task 2 collapsed and showed little task structure, indicating use of a distinct subspace. **D** Manifold separation predicts re-learning performance, similar to the experimental data (Kim et al., 2025).

We next asked how task-relevant representations were organized within these manifolds. For each task, we identified a choice axis separating left and right responses and examined the contribution of individual neurons to these axes (Methods). Visualization of neuronal weight contributions revealed that the choice axes for Task 1 and Task 2 were approximately orthogonal (Fig. 3B), indicating that the two stimulus–response mappings were embedded along distinct neural dimensions. In contrast, such orthogonal representations were not observed when the network was trained using FB1 throughout learning (Supplementary Fig. 5B).

To further characterize manifold separation, we defined a task-specific subspace by performing PCA on Task 1 population trajectories and projected activity from all training phases into this subspace (see Methods). We confirmed that the first three principal components accounted for more than 95% of the variance in Task 1 activity. Within this Task 1 subspace, left and right trajectories were clearly separated during Task 1 learning and re-learning, whereas trajectories from Task 2 collapsed and showed little task structure (Fig. 3C). This indicates that Task 2 activity occupied a different subspace and did not interfere with the previously learned Task 1 manifold. When feedback switching was omitted, trajectories from both tasks occupied the same subspace and the choice axis was preserved across conditions (Supplementary Fig. 5A), confirming that feedback switching is critical for manifold separation.

Finally, we asked whether the degree of manifold separation quantitatively predicts re-learning speed. Previous experimental work has shown that the angle between choice axes across tasks correlates with re-learning performance (Kim et al., 2025). To test whether the same relationship holds in our model, we systematically varied the alignment between choice axes across contexts and measured Task 1 re-learning speed (Methods). Both in the model and in experimental data, re-learning was fastest when choice axes were near-orthogonal and progressively slower as the axes became more overlapping (Fig. 3D). These results demonstrate a direct link between manifold separation of task representations and behavioral measures of interference.

In summary, switching feedback pathways reorganizes population activity into task-specific, low-dimensional manifolds. These manifolds are characterized by orthogonal choice axes across tasks and are selectively engaged during learning and re-learning, providing a geometric substrate for reducing interference between tasks.

### Autonomous emergence of task-specific feedback without external switching

So far, for the sake of simplicity, feedback signals were preconfigured and switched externally. We next asked whether such operations could instead be implemented autonomously. To test this possibility without relying on explicit task labels, we introduced a state-dependent feedback generation mechanism. Here, we assumed that feedback pathways were not manually assigned to tasks but were generated based on the current synaptic weight structures. Specifically, the feedback vector was updated with an element-wise product between readout weights weighted by readout activity and input weights weighted by the current input (Supplementary Fig. 6A). As learning progressed, the autonomously generated feedback vectors gradually diverged into distinct task-specific structures. During Task 2 learning, feedback became increasingly dissimilar to the Task 1 feedback vector, whereas during Task 1 re-learning it was progressively reinstated (Supplementary Fig. 6B, C). This divergence of feedback signals selectively stabilized task-specific activity patterns, allowing different neural representations to emerge purely through self-organized dynamics, without any external task cue. For clarity and consistency, all subsequent analyses use externally switched feedback signals, allowing us to isolate the role of feedback geometry without additional variability introduced by autonomous feedback generation.

### Orthogonal task manifolds suppress interference through geometric isolation

To understand the underlying mechanism by which interference is suppressed during continual learning in the proposed model, we examined the effect of learning-induced changes in recurrent connectivity during Task 2 on the dynamics of previously learned Task 1. Figure 4A illustrates the conceptual difference between orthogonal and overlapping manifold organization. We hypothesized that in the orthogonal case, synaptic updates induced during Task 2 learning minimally perturb population trajectories embedded in the Task 1 manifold, resulting in low interference. In contrast, when task representations occupy shared manifolds, identical synaptic updates substantially distort Task 1 trajectories, leading to high interference.

**Figure 4.**
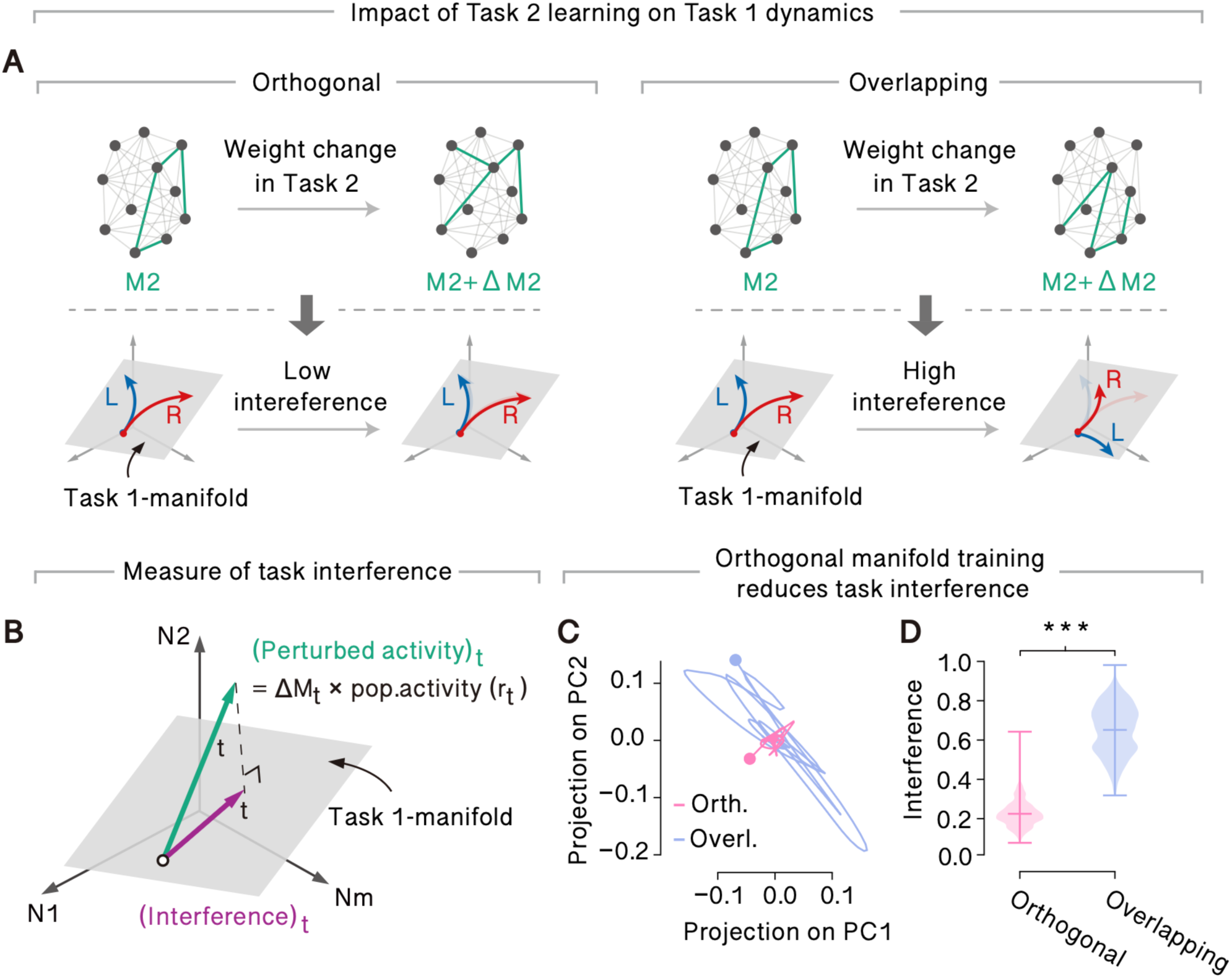
Orthogonal task manifolds minimize interference between tasks. **A** Conceptual schematic illustrating how learning-induced synaptic changes during Task 2 affect previously learned Task 1 dynamics under orthogonal versus overlapping manifold organization. In the orthogonal case (left), recurrent weight updates induced during Task 2 learning minimally perturb population trajectories embedded in the Task 1 manifold, resulting in low interference. In contrast, when task representations occupy overlapping manifolds (right), the same synaptic updates substantially distort Task 1 trajectories, leading to high interference. **B** Measure of task interference by projection of learning-induced activity perturbations onto the Task 1 manifold. During Task 2 learning, the instantaneous activity perturbation was computed at each learning step and projected onto the low-dimensional subspace defined by the principal components of the Task 1 manifold. The magnitude of this projection provides a measure of how strongly learning in Task 2 drives activity along previously learned task-relevant directions. **C** Example trajectories of projected perturbations within the Task 1 subspace. **D** Time-averaged magnitude of the projected perturbations shown in C. In D, P values were obtained from two-sided Welch’s t-tests across 20 independent simulations (***P < 0.001).

To test this hypothesis, we measured how learning-induced activity perturbations during Task 2 projected onto the Task 1 manifold. Specifically, at each learning step we computed the instantaneous activity perturbation during Task 2 learning and projected this vector onto the Task 1 manifold (Fig. 4B). This projection quantifies the extent to which learning in Task 2 drives population activity along task-relevant directions previously used for Task 1, providing a direct measure of interference between tasks.

Consistent with our hypothesis, projections of the perturbed activity onto the Task 1 subspace were substantially larger in the overlapping manifold condition than in the orthogonal manifold condition. In the overlapping case, projected trajectories spanned a broad region of the Task 1 manifold, whereas in the orthogonal case they remained tightly confined (Fig. 4C). Averaging the magnitude of these projections over time revealed significantly stronger interference in the overlapping condition compared to the orthogonal condition (Fig. 4D; ***P < 0.001, two-sided t-test).

Altogether, these results demonstrate that orthogonal manifold organization suppresses task interference by geometrically isolating learning-induced perturbations away from previously learned task subspaces.

### Task-specific memories are encoded in distinct connectivity modes

The results so far demonstrate that orthogonal feedback structures promote the formation of distinct task-specific manifolds avoiding interference. We next asked whether these task memories are stored in distinct synaptic connectivity patterns within the recurrent network.

To address this, we analyzed the recurrent weight matrix after Task 1 learning using singular value decomposition (SVD). The leading two singular modes captured the dominant connectivity structure underlying Task 1 performance (Fig. 5A), corresponding to the binary choice mappings (left and right) (Methods). We refer to this low-dimensional connectivity structure as engram 1. Similarly, after subsequent learning of Task 2 with orthogonal feedback FB2, we performed SVD on the updated recurrent weight matrix. Four dominant singular modes emerged, reflecting the combined representation of both tasks (Fig. 5A). Two of these modes closely matched the original Task 1 engram, while the remaining two captured novel connectivity patterns specific to Task 2. We therefore defined these additional modes as new connectivity modes.

**Figure 5.**
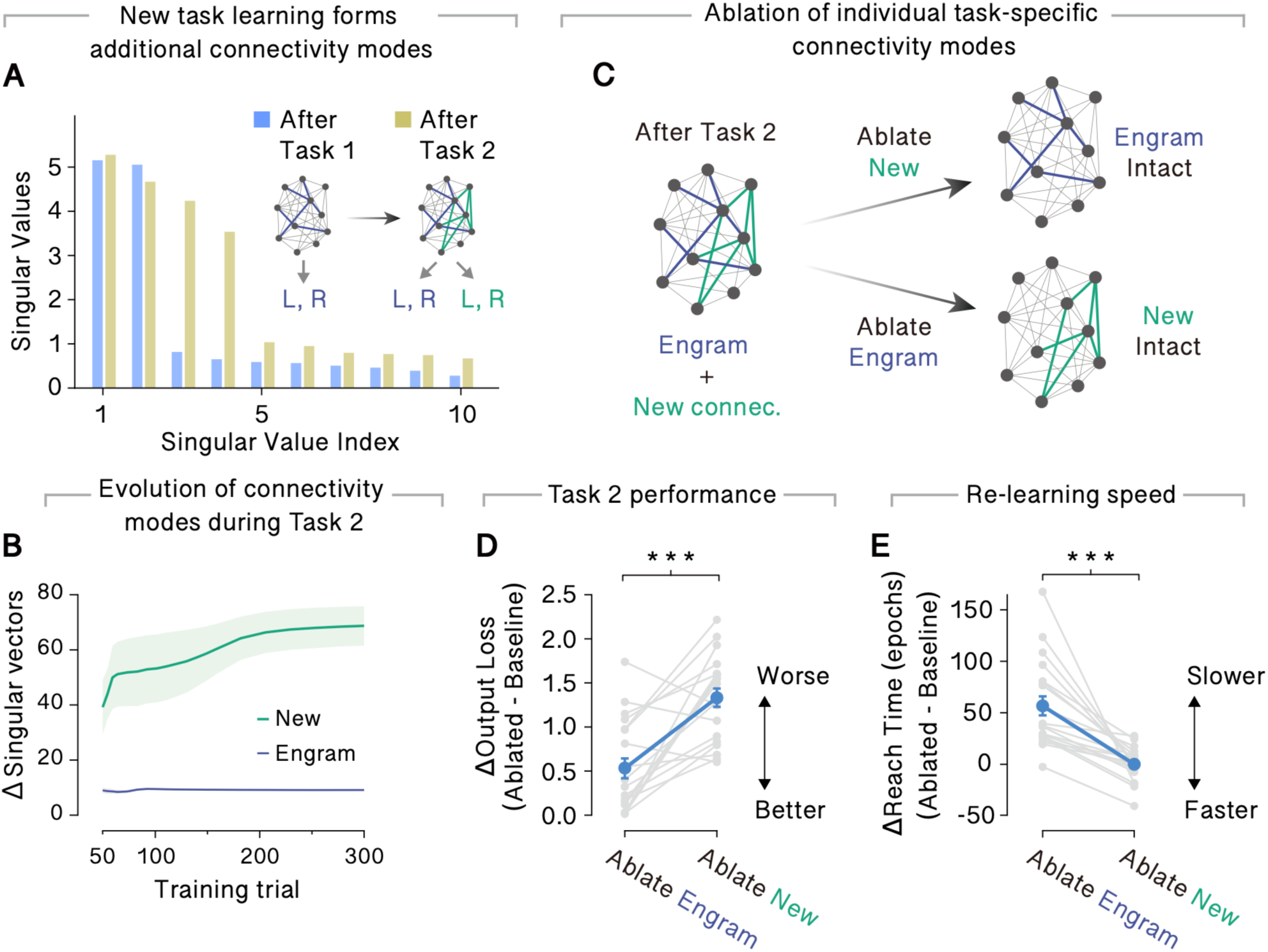
Task memories are encoded in separable connectivity modes. **A** Singular value decomposition (SVD) of the recurrent weight matrix after Task 1 and Task 2 learning. After Task 1 learning, two dominant singular modes emerge, corresponding to the binary choice mappings. After subsequent Task 2 learning with orthogonal feedback, four dominant modes are present, reflecting the combined representation of both tasks. **B** Temporal evolution of connectivity modes during Task 2 learning. Shaded areas represent s.d. over 5 independent simulations. **C** Schematic illustration of synaptic connectivity organization. After Task 2 learning, additional connectivity modes emerge in orthogonal directions while the original engram is preserved (left). Selective ablation of either the Task 1 engram or the newly formed connectivity modes (right). **D** Effects of selective connectivity ablation on Task 2 performance are shown. **E** Effects of selective connectivity ablation on Task 1 re-learning. In D and E, P values were obtained from two-sided Welch’s t-tests across 20 independent simulations (***P < 0.001).

The above results suggest that synaptic connectivity encoding Task 1 remains preserved during subsequent Task 2 learning, while new task-specific connectivity is added along orthogonal directions. To directly test this possibility, we examined how these connectivity modes evolved over the course of Task 2 learning. To this end, we tracked the similarity between the singular vectors defining engram 1 at the end of Task 1 learning and the corresponding singular vectors during Task 2 training (Methods). Strikingly, engram 1 remained highly stable throughout Task 2 learning, whereas the new connectivity modes progressively diverged as Task 2 learning proceeded (Fig. 5B). This indicates that Task 2 learning primarily recruited new, distinct connectivity modes while preserving the synaptic structure encoding Task 1.

We next tested whether these two connectivity modes were functionally necessary for task performance. After Task 2 learning, we selectively ablated either engram 1 or the newly formed connectivity modes and examined their effects on both Task 2 performance and Task 1 re-learning (Fig. 5C). Ablation of the newly formed connectivity modes significantly impaired Task 2 performance compared to ablation of engram 1, indicating that the new modes are essential for expressing Task 2 computations (Fig. 5D). In contrast, ablation of engram 1 significantly slowed re-learning relative to ablation of the new connectivity modes (Fig. 5E).

Together, these results provide direct causal evidence that orthogonal feedback structures give rise to separable synaptic connectivity modes within recurrent networks. Task memories are embedded as distinct low-rank connectivity modes within the recurrent weight matrix, enabling multiple tasks to coexist without mutual interference and accounting for the feedback-dependent re-learning dynamics observed earlier.

### Continual learning of high-dimensional naturalistic stimuli

To test whether orthogonal manifold organization generalizes beyond simple binary choice tasks, we trained networks to learn and replay high-dimensional natural movie sequences. Two distinct video clips (Movie 1 and Movie 2), each consisting of 150 frames, served as target outputs, with each frame corresponding to 1 ms. Each frame comprised 270 × 270 pixels with three color channels (RGB), yielding 218,700-dimensional output targets, far exceeding the 500 recurrent units in the network (Fig. 6A; Supplementary Fig. 7). At the beginning of each trial, network activity was initialized to predefined states drawn from a zero-mean unit-variance Gaussian distribution, which were held fixed across learning and testing. Importantly, the same set of initial network states was used for both Movie 1 and Movie 2, isolating the effect of feedback-driven reorganization of recurrent dynamics.

**Figure 6.**
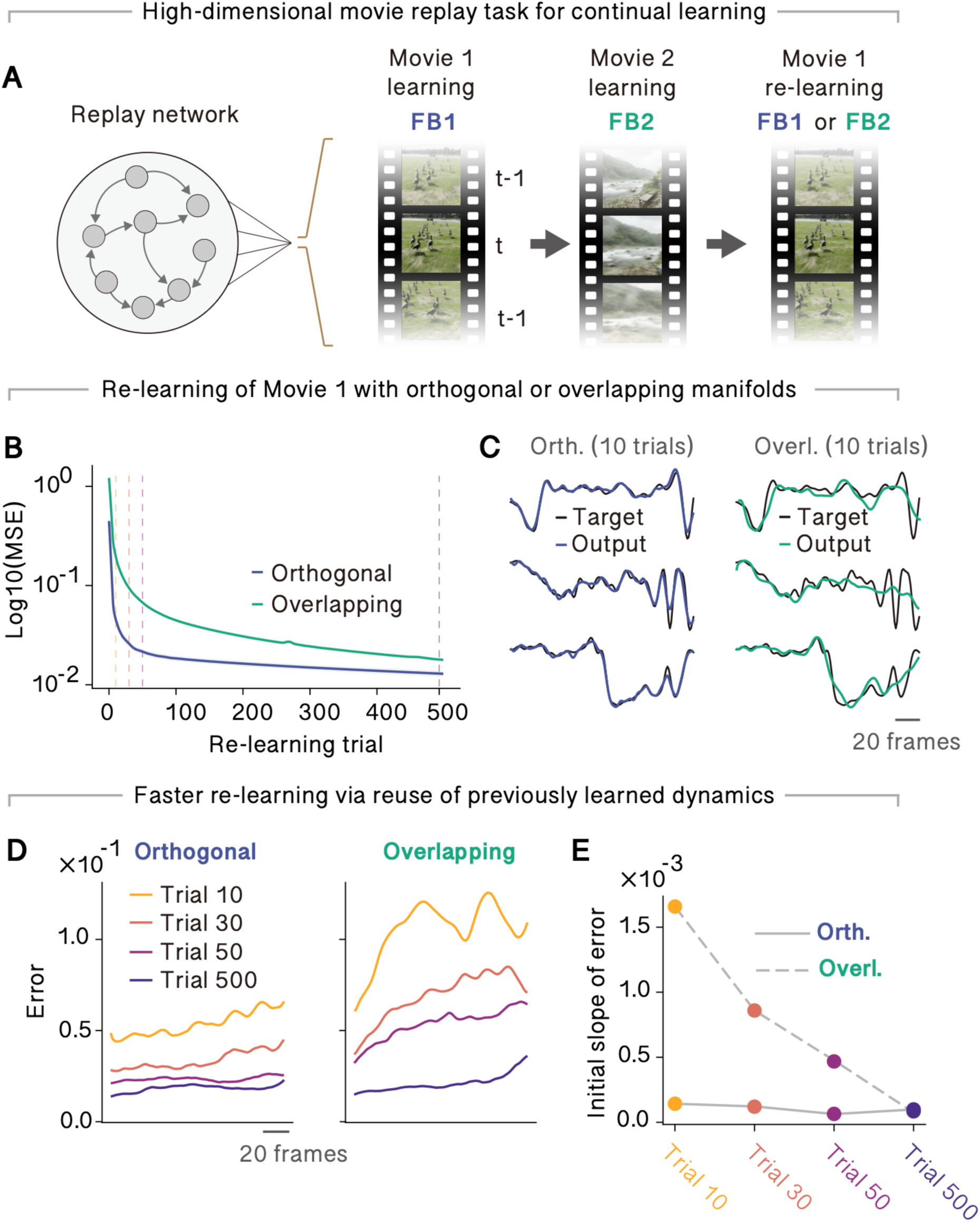
Feedback-driven orthogonal manifolds enable continual learning of high-dimensional naturalistic stimuli. **A** Schematic of the natural movie replay task and training protocol. Two high-dimensional natural video sequences (Movie 1 and Movie 2) were sequentially learned using distinct feedback pathways (FB1 and FB2). **B** Training error during re-learning of Movie 1 under orthogonal (FB1) and overlapping (FB2) conditions. **C** Example pixel target traces and corresponding learned readout dynamics after 10 re-learning trials under aligned and misaligned feedback conditions. **D** Frame-wise replay error across time shown for multiple re-learning trials. **E** Slopes of the frame-wise replay error curves shown in D.

As in previous experiments, Movie 1 was first learned using feedback FB1, followed by learning of Movie 2 using FB2. The network was then retrained on Movie 1 under either the aligned feedback FB1 (orthogonal) or the misaligned feedback FB2 (overlapping) (Fig. 6A). Notably, unlike the binary choice task, no external input was provided during movie replay, and successful learning required the network to generate autonomous dynamics rather than converge to fixed points. Despite the large mismatch between target dimensionality and network size, networks in both conditions successfully re-learned Movie 1, with training errors converging to near-zero values (Fig. 6B; Supplementary Fig. 8). However, re-learning speed showed a strong dependence on feedback identity. Re-learning with FB1 was substantially faster than re-learning with FB2 (Fig. 6B, C), indicating that the Movie 1 dynamics were preserved during Movie 2 learning when feedback-specific manifolds were orthogonalized.

To examine how network dynamics were reorganized during re-learning, we analyzed frame-wise readout errors at multiple re-learning trials. Under the orthogonal condition, replay errors remained low and approximately uniform across frames even at early re-learning stages, and decreased further with training (Fig. 6D). In contrast, under the overlapping condition, replay errors increased progressively across frames during early re-learning, indicating cumulative temporal drift, and only gradually improved with extensive re-learning (Fig. 6E). These distinct error profiles suggest that, in the orthogonal condition, re-learning reused preserved Movie 1 trajectories, whereas re-learning in the overlapping condition required reconstruction of the movie dynamics from scratch.

Together, these results demonstrate that feedback-driven orthogonal manifold organization extends to complex, high-dimensional temporal patterns. By linking distinct plasticity feedback pathways to different spatiotemporal target patterns, recurrent networks minimize memory interference and enable efficient continual learning, even for high-dimensional naturalistic inputs.

## Discussion

We demonstrated that recurrent neural networks can continually acquire multiple tasks while minimizing catastrophic interference by organizing population activity into distinct, orthogonal manifolds. Inspired by predictive learning mechanisms (Urbanczik, R., & Senn, 2014; Asabuki & Fukai, 2020; Recanatesi et al., 2021; Levenstein et al., 2024), we showed that task-specific feedback signals guide predictive synaptic plasticity to learn separable subspaces of recurrent dynamics, enabling rapid switching between tasks while preserving previously learned computations. Importantly, previously learned task representations are not erased but preserved as latent dynamical structures that can be rapidly reactivated through appropriate re-learning. This principle generalizes from simple decision-making tasks to high-dimensional naturalistic stimuli, where distinct feedback pathways preserve latent autonomous dynamics and enable efficient re-learning of complex movie sequences. Together, these results propose feedback-driven manifold switching as a biologically plausible mechanism for continual learning in recurrent circuits.

From a theoretical perspective, our work complements and extends prior proposals that memory stability can be achieved by allocating information across orthogonal or minimally overlapping representational dimensions (Fusi & Abbott, 2007; Benna & Fusi, 2016; Duncker et al., 2020). While previous models often enforced such separation through architectural constraints or global optimization objectives, we show that orthogonal organization can instead emerge incrementally through local, feedback-driven predictive plasticity. Crucially, this organization can arise without explicit task labels provided to recurrent units during task execution. Rather, feedback geometry biases synaptic updates toward task-aligned modes, shaping both population activity and synaptic structure.

Our results also resonate with a growing body of work suggesting that, while memory is stored in synaptic modifications, its functional expression in recurrent circuits is inherently dynamical. Theoretical studies have shown that stable computation can be supported by attractors, low-dimensional trajectories, or slow modes embedded in recurrent dynamics (Barak & Tsodyks, 2007; Mongillo et al., 2008; Chaudhuri & Fiete, 2016). Within this framework, learning inevitably reshapes the geometry of the dynamical landscape through synaptic plasticity. Crucially, our proposed learning mechanism induces orthogonal task-specific manifolds within this landscape. This organization explains why task-aligned perturbations selectively impair corresponding behaviors while sparing others, and why the degree of geometric alignment predicts re-learning speed.

At the circuit level, our findings suggest a close relationship between feedback signals and the organization of task representations. Cortical feedback pathways are known to carry contextual, predictive, and goal-related information and to exert powerful modulatory effects on local computations (Mumford, 1992; Rao & Ballard, 1999; Bastos et al., 2012; Zhang et al., 2014; Keller & Mrsic-Flogel, 2018). Recent experimental work has highlighted the role of feedback and apical dendritic input in gating plasticity and shaping learning-related changes in cortical circuits (Larkum, 2013; Phillips et al., 2016). In this light, feedback-driven manifold switching provides a concrete computational mechanism through which top-down signals could route learning into task-specific dynamical subspaces (Fişek et al., 2023), enabling flexibility without catastrophic interference.

Our results further connect to experimental observations of representational drift, where population-level codes remain functionally stable despite substantial turnover in single-neuron tuning (Driscoll et al., 2017; Rule et al., 2019; Driscoll et al., 2022). We show that such apparent stability can arise naturally when task-specific computations are embedded in low-dimensional manifolds that are preserved across learning. Even as synaptic weights change, the underlying task-relevant subspaces remain accessible, allowing rapid reinstatement of behaviorally relevant dynamics. This view aligns with experimental findings that learning and context switches often correspond to rotations or reconfigurations of population activity within a stable latent space rather than complete remapping (Stavisky et al., 2017; Gallego et al., 2018; Vyas et al., 2020; Gallego et al., 2020).

The persistence of task-specific synaptic connectivity modes in our model also parallels emerging concepts from the memory engram literature. Experimental studies have shown that memories are stored in distributed but selectively reactivatable neural ensembles and circuits, which can persist in a latent state and be reinstated by appropriate cues (Tonegawa et al., 2015; Kitamura et al., 2017; Josselyn & Tonegawa, 2020). Our results suggest that such engrams may consist of structured synaptic modifications that collectively reshape the geometry of recurrent dynamics to support population activity (Peron et al., 2020), which, in our framework, is organized within orthogonal manifolds. Disrupting such synaptic connectivity selectively impairs associated behaviors, consistent with causal manipulations of engram circuits observed in vivo.

Several open questions follow from our work. First, while we demonstrated orthogonal organization for pairs of tasks, scaling this mechanism to many interleaved tasks raises questions about capacity limits and how networks balance memory specificity with generalization. Partial overlap between manifolds may support transfer learning and compositionality but could also reintroduce interference. Second, our model assumes the availability of task-specific feedback signals. In biological systems, such feedback may be learned, inferred, or contextually gated, suggesting future work on how feedback routing itself is acquired. Third, extending these principles to spiking networks with additional biophysical constraints, including Dale’s law and spike-timing-dependent plasticity, will be important for closer links to neural circuitry.

Our framework generates several experimentally testable predictions. Continual learning should be accompanied by rotations of population activity into task-specific subspaces that can be tracked using dimensionality reduction techniques. Direct perturbation of task-specific feedback pathways is predicted, in principle, to disrupt manifold separation and increase interference between tasks. More indirectly, trial-to-trial fluctuations in feedback-related activity should correlate with reduced subspace separation and impaired behavioral performance. Such predictions can be tested in rodents or primates performing context-dependent tasks engaging recurrent cortical networks.

Finally, our findings have implications for artificial intelligence. Most continual learning algorithms focus on stabilizing synaptic parameters, replaying past experiences, or expanding network capacity. Our results highlight an alternative principle: controlling the geometry of internal representations. Learning rules, architectural biases, or feedback mechanisms that promote manifold separation may enable flexible continual learning without freezing plasticity. More broadly, the strategic organization and reallocation of representational geometry may constitute a general design principle shared by biological and artificial learning systems.

In conclusion, we show that feedback-driven predictive plasticity provides a simple and biologically plausible mechanism for organizing recurrent dynamics into task-specific manifolds, enabling continual learning without catastrophic interference. By shaping the geometry of population activity rather than protecting individual synapses, recurrent circuits can flexibly acquire new skills while preserving previously learned computations. These results place feedback geometry at the center of the link between synaptic plasticity, population dynamics, and memory stability.

## Methods

### The neural network model

Our network consists of recurrently connected *N* rate-based units. The dynamics of membrane potential with the external input ***I*** was governed by the following equation:

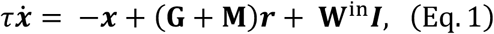

where the variable ***x*** is a membrane potential and ***r*** is a firing rate of each unit, defined as:

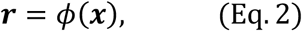

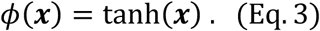

Here, we considered two types of recurrent connections: a strong, fixed connectivity **G** as well as initially weak and plastic connectivity **M** (Asabuki & Clopath, 2025). The strength of each connection was generated by a Gaussian distribution, and unless otherwise specified, with zero-mean and the standard deviation of 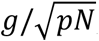, with (*p*, *g*) = (0.1, 1.2) for **G** and (*p*, *g*) = (1.0, 0.5) for **M**. The parameter *g* controls the gain of recurrent coupling and induces chaotic spontaneous activity when *g* > 1 (Sompolinsky et al., 1988). Although an additional plastic connectivity **M** was present, the fixed recurrent connectivity **G** provided the initial chaotic spontaneous activity of the network (Asabuki & Clopath, 2025). The matrix **W**^in^ is a feedforward connectivity projecting from the input layer, with each element independently drawn from a uniform distribution between −2 and 2, which is assumed to be fixed during the whole simulation. Note that external inputs were included only in the contingency reversal task. Details of the input implementation are described below. The parameter *τ* is a membrane time constant, which we set as *τ* = 50 (ms). The recurrent units project to readout units, of which the output value was defined as a linear summation of firing rates:

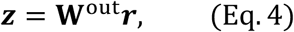

where **W**^out^ is a readout weight matrix, generated by a Gaussian distribution with zero mean and variance 1/N.

In all simulations, network activity was initialized at the beginning of each trial to a predefined state. This state was sampled once from a zero-mean unit-variance Gaussian distribution and was held fixed across all learning and testing.

To train the recurrent connections, we defined the feedback matrix **Q**, of which the elements were drawn from a uniform distribution. In the contingency reversal task, each element of **Q** was sampled from the uniform distribution between −3 and 3. In contrast, in the movie replay task in Figure 6, the elements were drawn from a uniform distribution between 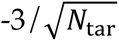 and 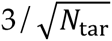, where *N*_tar_ is the number of target (teacher) signals. With the given feedback matrix **Q** for the recurrent plasticity, we trained the recurrent connection **M** by the following plasticity rule (Asabuki and Clopath, 2025):

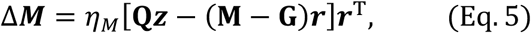

where T is a transpose. Note that the matrix **Q** does not appear in the neural dynamics (Eq. 1). Readout weights were trained by simple least-mean-square (LMS):

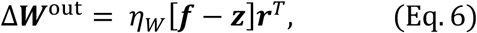

where ***f*** is a vector of teaching signal for the model’s outputs. Here, we used identical learning rates for both parameters, setting *η*_*M*_ = *η_W_* = 10^−4^.

### Contingency Reversal Task

The contingency reversal task consisted of two external input units and two output units. The mapping between inputs and outputs differed between Task 1 and Task 2, as described in the Results. Each trial lasted 3.5 s and was divided into two phases. During the first 1500 ms (hold period), one of the two input units was selected and set to *I*(*t*) = 1, while the non-selected input unit was held at zero throughout the trial. During this period, both output units were assigned a target value of −1. During the subsequent 2,000 ms (reaching phase), the hold input smoothly and rapidly decayed to zero. Concurrently, the output unit corresponding to the selected input transitioned from −1 to +1, whereas the non-corresponding output unit remained fixed at −1. This input–output contingency was reversed between Task 1 and Task 2.

### Measuring alignment between feedback and neural manifolds

In Figure 3A, we performed the following analysis at each training trial during the contingency reversal task. First, at each trial, we collected the network activity ***r***(t) ∈ ℝ^N^ across time and organized it into a matrix:

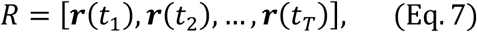

where *T* is the length of each trial. We then applied principal component analysis (PCA) to *R* to extract the dominant PCA modes of each task manifold, focusing on the first two principal components (PC1 and PC2), which were normalized to have unit length.

We performed the following analysis over two task-specific feedback matrices (**Q**_1_ and **Q**_2_). We explain in a case **Q** = **Q**_1_, where the columns represent the target directions:

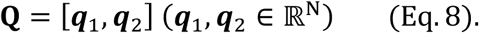

Each column vector ***q***_*i*_(*i* = 1,2) was also normalized to unit length prior to comparison. We then calculated the absolute value of cosine similarity between each ***q***_*i*_and each principal component vector (**PC**):

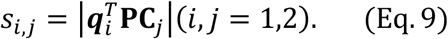

Since we have two possibilities over coupling of each ***q***_*i*_ and each **PC**, we computed

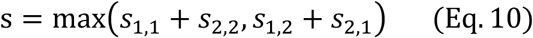

to record the individual cosine similarity scores for the selected optimal pairing in each trial. We compared the trajectories of such cosine similarity between feedback and **PC**s over the case **Q** = **Q**_1_ and **Q** = **Q**_2_.

### Visualization of weights on choice axis

In Figure 3B, for each task, we first defined a choice axis that captured population-level activity differences between left and right choices. Specifically, we collected the network activity vectors at the final time step of each trial, separately for trials in which the network chose the left or right option. Let ***r***_left_ and ***r***_right_ denote the average population activity vectors across left-choice and right-choice trials, respectively. The choice axis was then defined as the difference vector

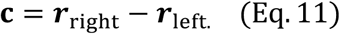

This procedure was applied independently to Task 1 and Task 2, yielding two choice axes **c**^(1)^ and **c**^(2)^, referred to as Choice-1 and Choice-2 in the Results section. Each axis is an N-dimensional vector, where N is the number of recurrent units in the network. For each neuron *i*, we then extracted the pair 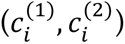, corresponding to its contribution to the Task 1 and Task 2 choice axes, respectively. The distribution of these neuron-wise contribution pairs was visualized as a two-dimensional histogram (Fig. 3B).

### Quantifying re-learning speed as a function of choice-axis alignment

In Figure 3D, to systematically control the alignment between task-specific choice axes, we generated the feedback vector for Task 2 as a linear combination of two orthogonal components. Specifically, we defined

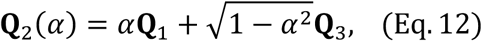

Where **Q**_1_ is the feedback vector used for Task 1, and **Q**_3_ is an auxiliary vector chosen to be orthogonal to **Q**_1_ direction. By varying the mixing coefficient α, we generated task contexts associated with different degrees of alignment between the resulting choice axes.

For each value of α, we computed the choice axes for Task 1 and Task 2 as described above and quantified their geometric overlap by calculating the normalized dot product between the two axes. We then measured the speed of Task 1 re-learning following Task 2 training under each condition. Re-learning speed was quantified as the number of trials required for the mean squared error (MSE) during Task 1 re-learning to fall below a fixed threshold of 10^−3^.

### Autonomous generation of feedback signals

In Supplementary Figure 6, at each time step, the feedback vector **Q**(t) was updated as

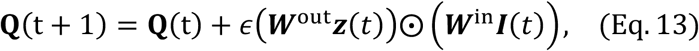

where ***W***^out^ denotes the readout weight vector, ***z***(*t*) is the readout activity, ***W***^in^ denotes the input weight vector, ***I***(*t*) is the current external input, and ⊙ indicates element-wise multiplication, and the learning rate was set to *∊* = 0.2. This operation generates a feedback signal that selectively reflects the conjunction of output-related and input-related synaptic pathways.

For clarity and consistency, all main-text analyses were performed using externally switched feedback signals, allowing us to isolate the role of feedback geometry without additional variability introduced by autonomous feedback generation.

### Analysis of interference between task manifolds

In Figure 4, to quantify interference between different manifolds, we evaluated how much the activity changes induced by synaptic updates during Task 2 were projected onto the feedback subspace associated with previously learned tasks (i.e., Task 1). At each learning step in Task 2, we defined the resulting change in internal activity, Δ***r***(t), as:

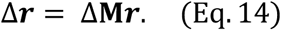

This approximates how the recurrent dynamics would be perturbed immediately after the synaptic update, based on the pre-update activity ***r***. Thus, Δ***r*** represents the change in neural activity caused by the synaptic plasticity event.

To measure interference, we projected the change in population activity Δ***r*** during learning of Task 2 onto the subspace spanned by the first two principal components of the population activity associated with Task 1. The projection was computed as

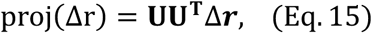

where the columns of **U** are the first two principal component vectors obtained from the population activity during Task 1, forming an orthonormal basis of the task-related subspace. The norm of the projected component was used to quantify the normalized interference magnitude:

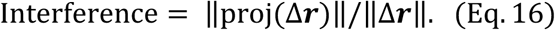

This quantity captures how much the synaptic updates during learning of a new task inadvertently perturb the previously formed neural manifold.

### Singular value decomposition of learned recurrent connectivity

In Figure 5A, to analyze how task memories are embedded in recurrent synaptic connectivity, we applied singular value decomposition (SVD) to the recurrent weight matrix **M** after learning. The matrix was decomposed as

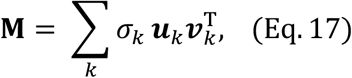

where *σ*_*k*_ are singular values, and ***u***_*k*_ and ***v***_*k*_ are the corresponding left and right singular vectors. Each rank-one term 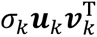 represents a structured connectivity mode that shapes population activity along the direction ***v***_*k*_ to downstream activity along ***u***_*k*_. In recurrent networks, such low-rank connectivity components selectively shape population dynamics along specific low-dimensional subspaces (Mastrogiuseppe & Ostojic, 2018). We therefore interpret dominant singular modes as connectivity structures that implement task-relevant computations.

After Task 1 learning, the first two singular modes accounted for the dominant low-rank structure of the recurrent weight matrix and corresponded to the binary choice mappings (left and right). We defined these two modes collectively as engram 1. After subsequent learning of Task 2 with orthogonal feedback, additional dominant singular modes emerged, reflecting newly formed task-specific connectivity. These modes were defined as new connectivity modes.

### Tracking stability of connectivity modes

In Figure 5B, to track the evolution of Task 1 connectivity during Task 2 learning, we identified the optimal correspondence between singular modes obtained after Task 1 learning and those obtained at each stage of Task 2 learning. Because singular vectors are not ordered across decompositions, we determined the matching that minimized the first principal angle between subspaces spanned by the corresponding left singular vectors ***u***_*k*_. Specifically, for each candidate pairing, we computed the first principal angle between the subspaces defined by the Task 1 singular vectors and the singular vectors at a given point during Task 2 learning and selected the combination that minimized this angle.

Using the matched singular vectors, we quantified the stability of each connectivity mode by computing the similarity between the Task 1 engram vectors at the end of Task 1 learning and their corresponding vectors during Task 2 training. In parallel, we tracked the evolution of newly formed connectivity modes once they became well-defined during Task 2 learning. Because switching to the Task 2 feedback initially induces transient and unstable network dynamics in which low-rank connectivity structure is not yet formed, connectivity mode tracking during Task 2 learning was initiated after the first 50 training trials.

### Simulation details

All simulations were performed in customized Python3 code written by Z.L. with numpy 1.26.4 and scipy 1.15.3. Differential equations were numerically integrated using an Euler method with integration time steps of 10 ms.

## Data availability

All numerical datasets necessary to replicate the results shown in this article can easily be generated by numerical simulations with the software code available upon publication. No datasets were generated during this study.

## Code availability

Example source codes used for the present numerical simulations and data analysis will be made available upon publication.

## Author Contributions

T.A. designed the study and wrote the paper. Z.L. performed numerical simulations. A.K. and Y.O. contributed to the design of the analyses. All authors discussed the results and contributed to editing the manuscript.

## Acknowledgments

This work was supported by RIKEN Center for Brain Science (TA) and RIKEN Cluster for Pioneering Research (TA). We are grateful to Jay Hennig for his fruitful discussions.

## Supplementary Figures

**Supplementary Figure 1:**
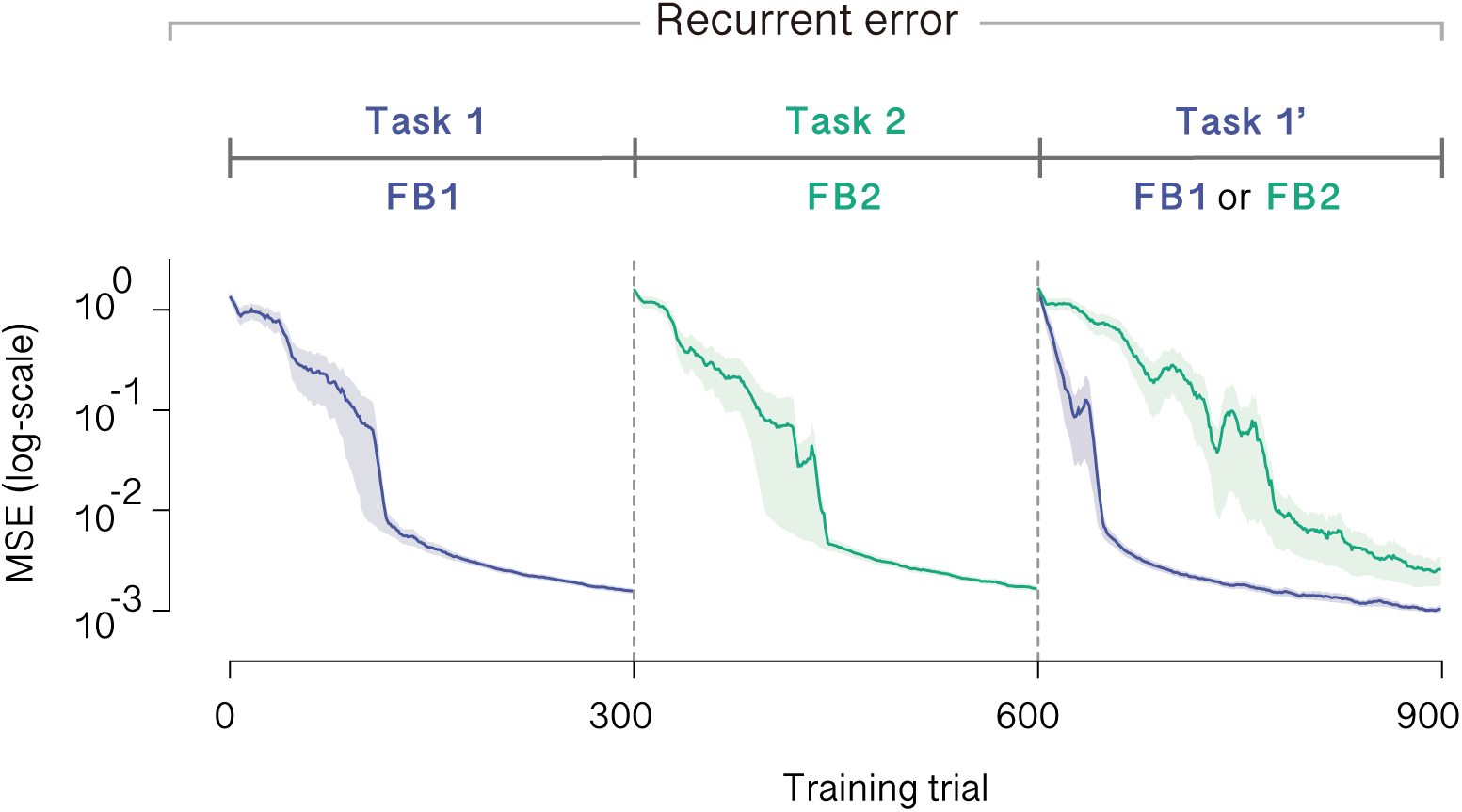
Faster relearning of recurrent dynamics with task-matched feedback. Learning curves of recurrent error during relearning for Task 1 under different feedback conditions are shown.

**Supplementary Figure 2:**
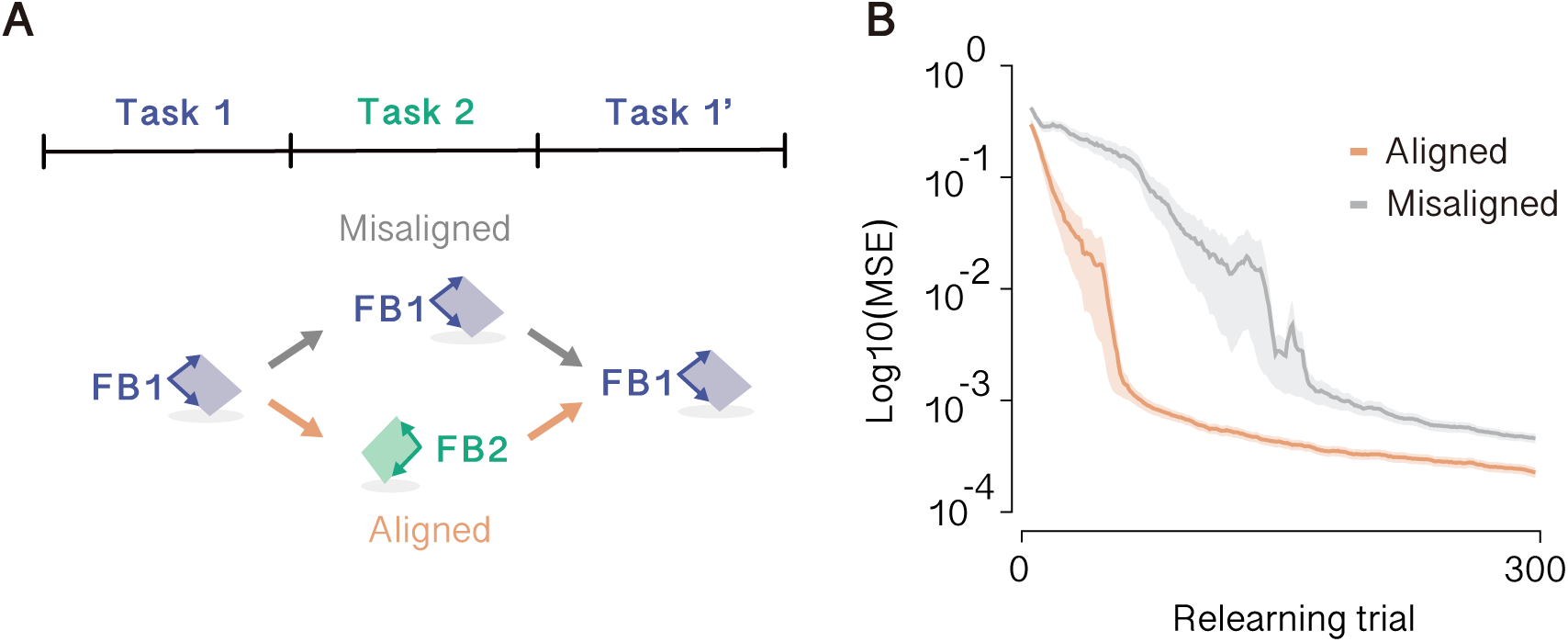
Task-dependent feedback signals determines the speed of relearning. **A.** Schematic of feedback conditions across task phases. During Task 2 learning, feedback was either aligned with the current task (FB2, aligned) or mismatched to the task (FB1, misaligned). **B.** Learning curves during Task 1 relearning under aligned and misaligned feedback conditions. When feedback during Task 2 learning was aligned to Task 2 (FB2), relearning of Task 1 proceeded rapidly.

**Supplementary Figure 3:**
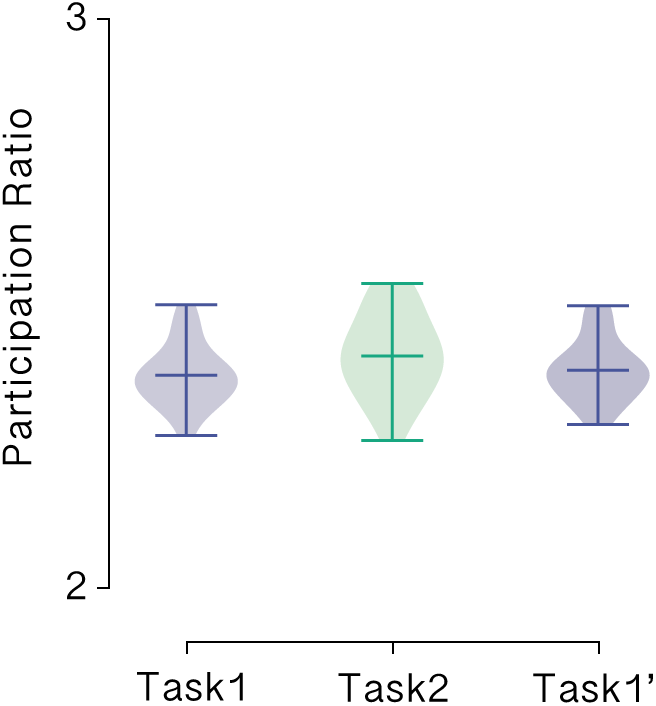
Stable dimensionality of network activity across learning. Participation ratio of population activity during Task 1 learning, Task 2 learning, and Task 1 relearning (Task 1′) under the aligned feedback condition. The participation ratio remained approximately constant across all phases, indicating that the dimensionality of network activity was preserved throughout learning and relearning. Each distribution summarizes values obtained from 20 independent simulation runs.

**Supplementary Figure 4:**
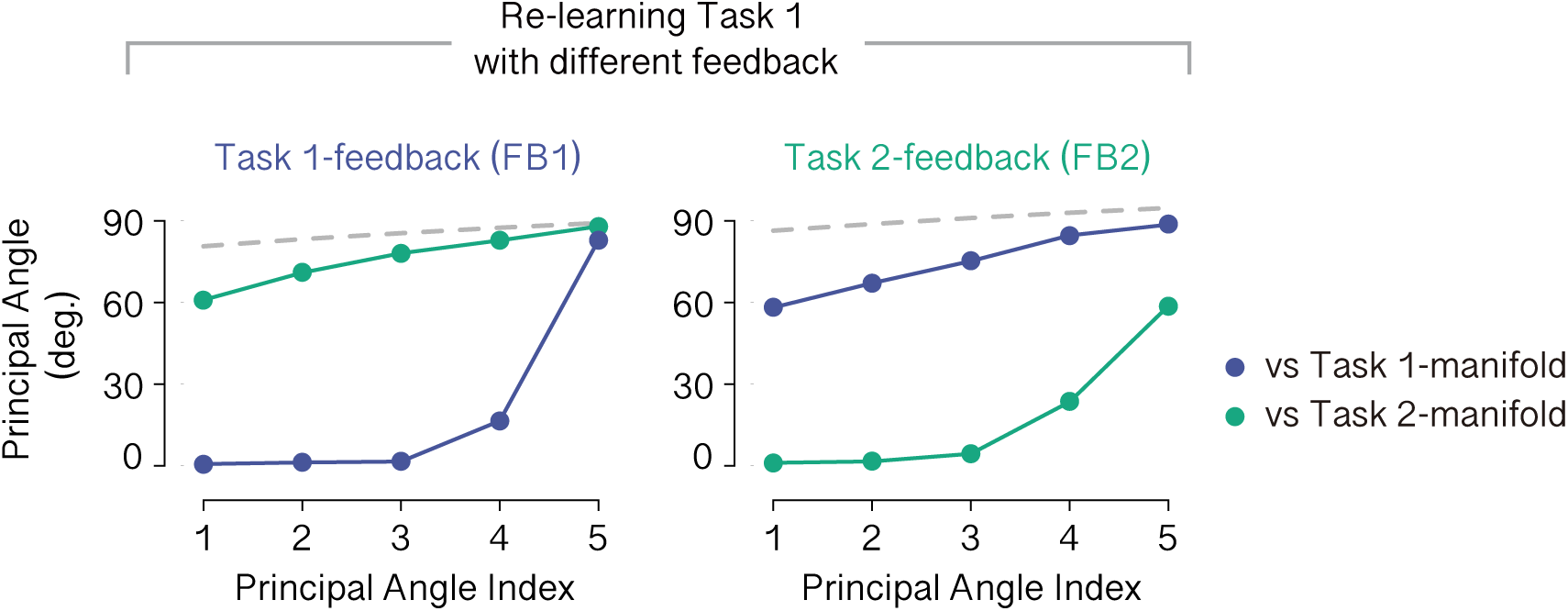
Feedback-dependent switching of task manifolds during re-learning. Principal angles between population activity during Task 1 re-learning and task-specific manifolds are shown for re-learning under Task 1 feedback (FB1) and Task 2 feedback (FB2). FB1 selectively aligned activity with the Task 1 manifold, whereas FB2 aligned activity with the Task 2 manifold, indicating feedback-dependent switching of low-dimensional task representations. Gray lines indicate chance levels.

**Supplementary Figure 5:**
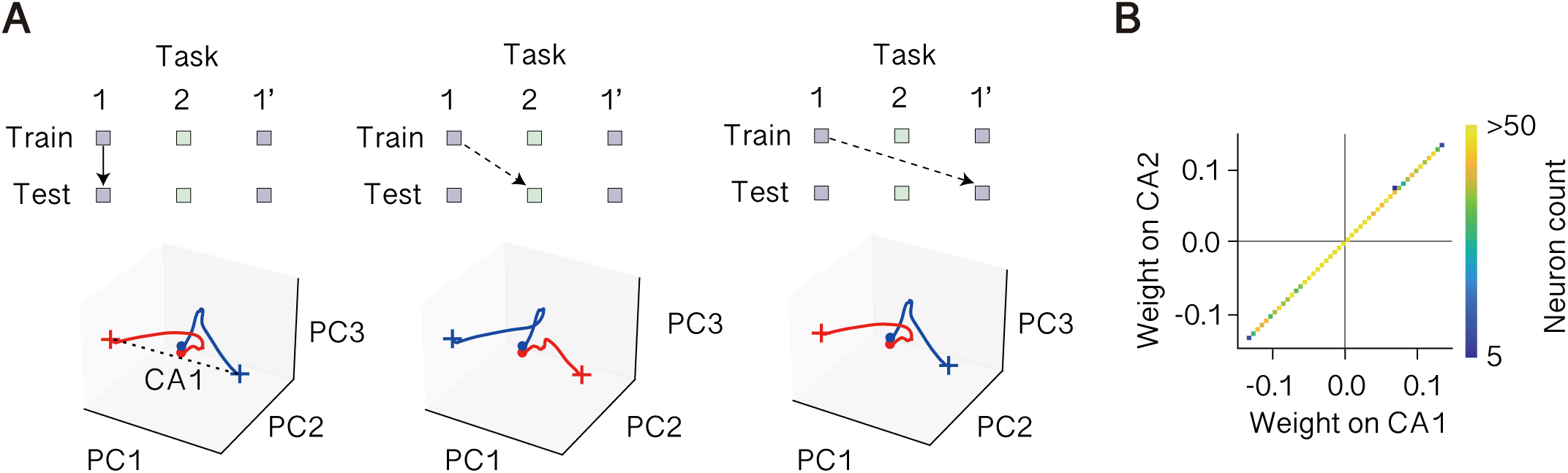
Constant feedback signals lead to no manifold switching. **A.** In the misaligned feedback condition, population activity projected onto all task-defined subspaces exhibited two trajectories corresponding to left and right choices, indicating that both tasks occupied overlapping subspaces. **B.** Choice axes separating left and right responses were aligned across Task 1 and Task 2 in the misaligned feedback condition.

**Supplementary Figure 6:**
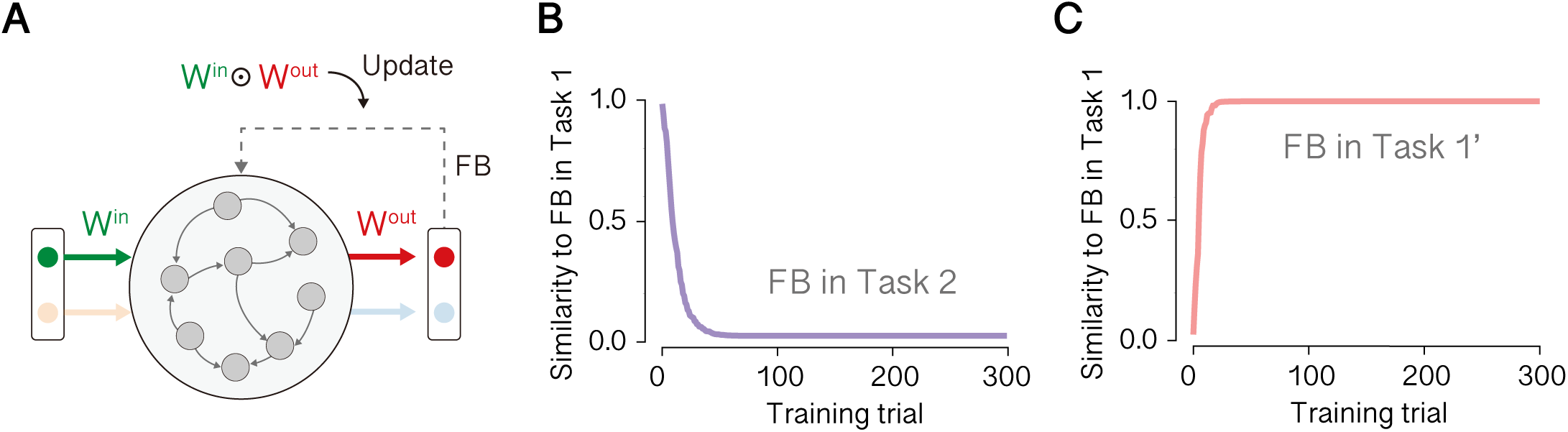
Autonomous feedback generation switches between task-specific feedback signals. **A.** The feedback vector was updated as a slow activity-dependent combination of input and readout pathways, implemented as a low-pass integration of their element-wise interaction. **B.** During Task 2 learning, the similarity between the autonomously generated feedback vector and the feedback vector characteristic of Task 1 progressively decreased, indicating a gradual divergence from the Task 1 feedback. **C.** During Task 1 re-learning, the similarity to the Task 1 feedback vector increased over time, reflecting reinstatement of the Task 1 feedback structure without any external task cue.

**Supplementary Figure 7:**
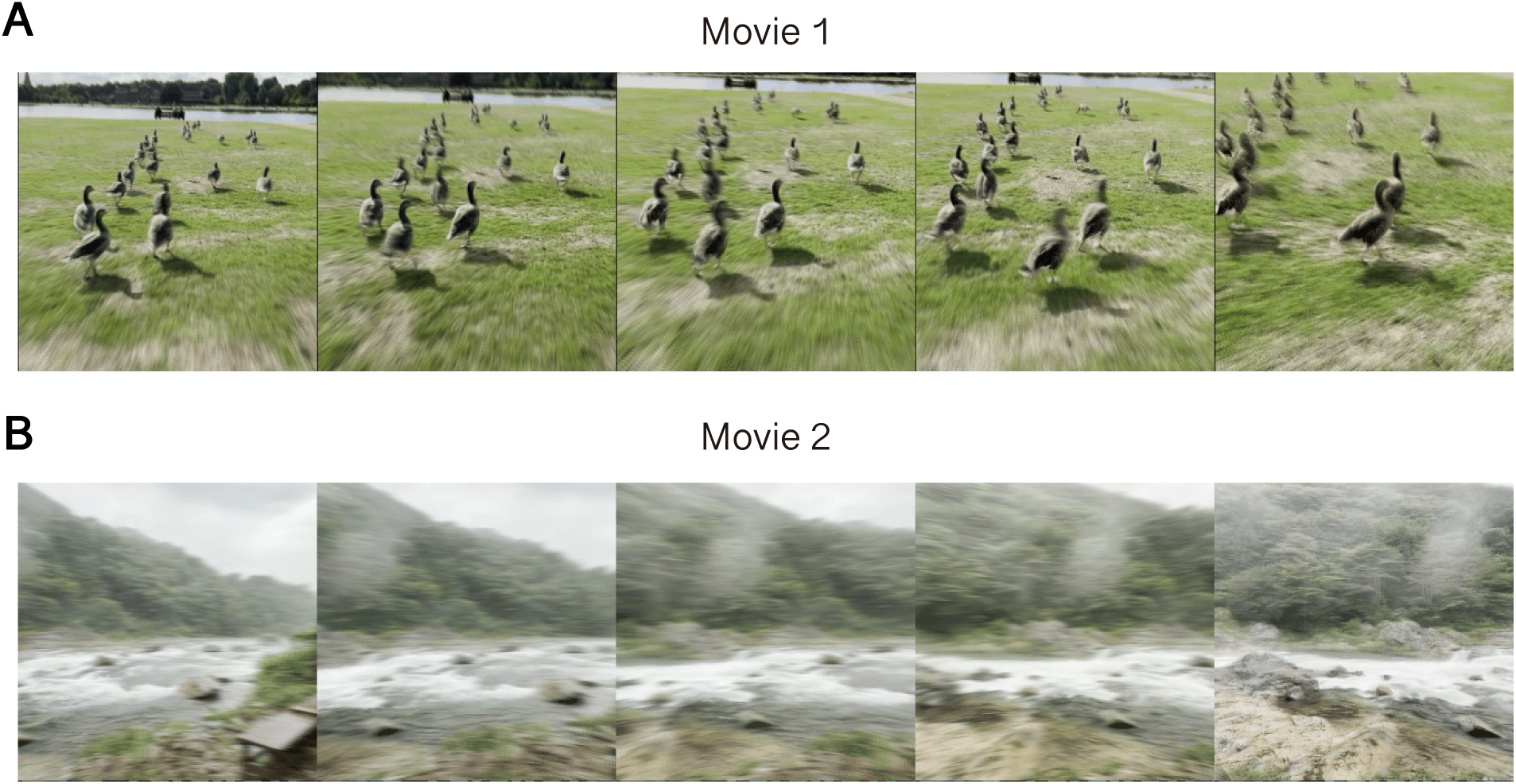
Example frames of teacher movies. **A.** Example frames from Movie 1 used as the teacher signal in the movie task. Five frames sampled from the sequence are shown. **B.** Same as A, but for Movie 2

**Supplementary Figure 8:**
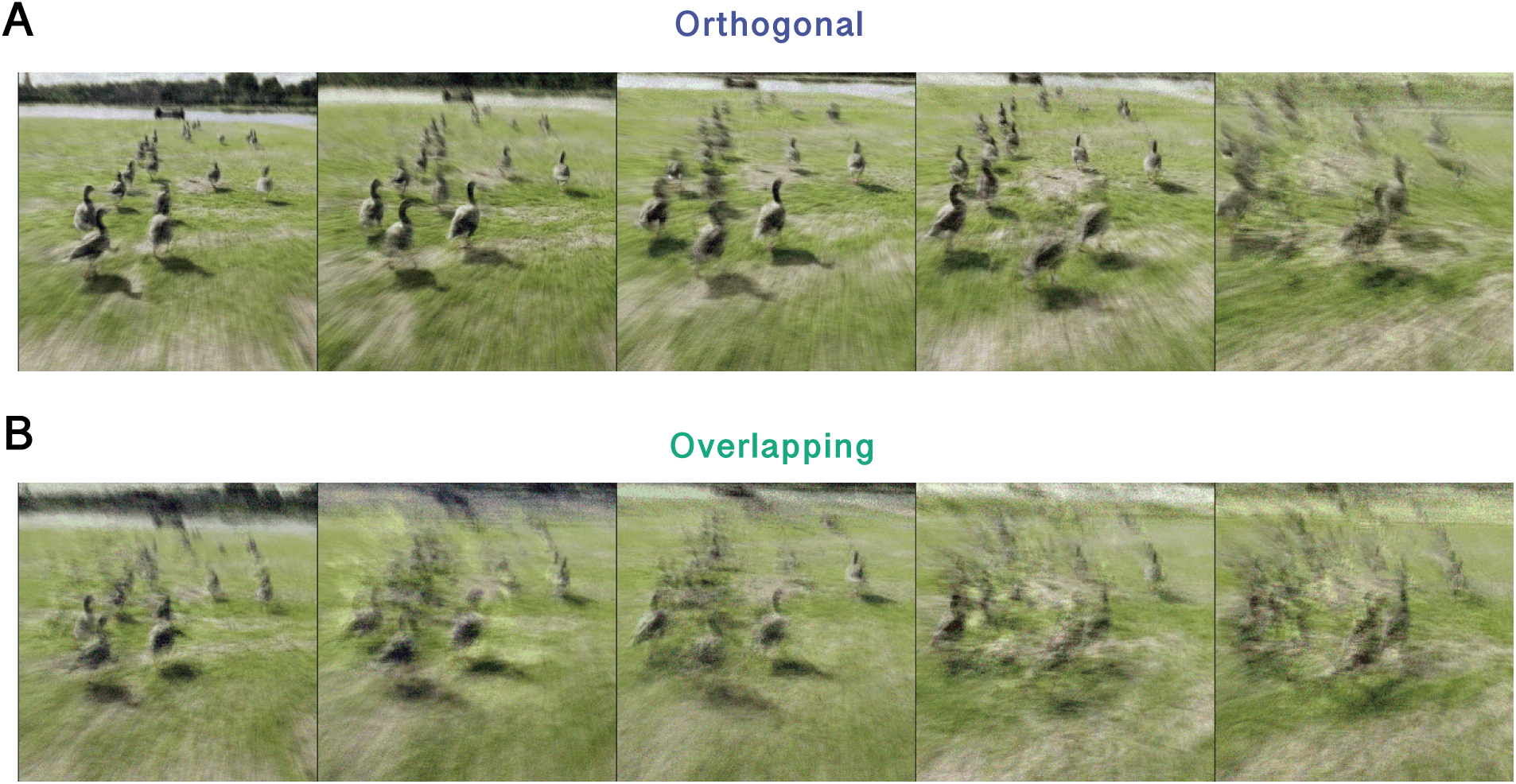
Example frames of replayed movies. **A.** Example frames from the replayed movie after relearning over 10 training trials in the aligned feedback condition. Five frames sampled from the sequence are shown. **B.** Same as A, but for the misaligned feedback condition.

